# Circular RNA expression in human hematopoietic cells is widespread and cell-type specific

**DOI:** 10.1101/300293

**Authors:** Benoit P Nicolet, Sander Engels, Francesca Aglialoro, Emile van den Akker, Marieke von Lindern, Monika C Wolkers

## Abstract

Hematopoietic stem cells differentiate into a broad range of specialized blood cells. This process is tightly regulated and depends on transcription factors, micro-RNAs, and long non-coding RNAs. Recently, also circular RNA (circRNA) were found to regulate cellular processes. Their expression pattern and their identity is however less well defined. Here, we provide the first comprehensive analysis of circRNA expression in human hematopoietic progenitors, and in differentiated lymphoid and myeloid cells. We here show that the expression of circRNA is cell-type specific, and increases upon maturation. circRNA splicing variants can also be cell-type specific. Furthermore, nucleated hematopoietic cells contain circRNA that have higher expression levels than the corresponding linear RNA. Enucleated blood cells, i.e. platelets and erythrocytes, were suggested to use RNA to maintain their function, respond to environmental factors or to transmit signals to other cells via microvesicles. Here we show that platelets and erythrocytes contain the highest number of circRNA of all hematopoietic cells, and that the type and numbers of circRNA changes during maturation. This cell-type specific expression pattern of circRNA in hematopoietic cells suggests a hithero unappreciated role in differentiation and cellular function.

## INTRODUCTION

Each day more than 10^12^ cells are produced in the bone marrow from hematopoietic stem cells (HSCs). HSCs differentiate into various progenitor cells, which in turn generate many types of myeloid and lymphoid cells (1). This process requires a tight regulation of gene expression. Transcription factors, long non-coding RNAs (lncRNAs), and microRNAs (miRNAs) contribute to differentiation (2–4). In addition to miRNA and lncRNA, also other non-coding RNAs emerge as important regulatory factors. Recently, it was shown that an alternative splicing mechanism can give rise to stable circular RNA (circRNA) with distinct regulatory capacity (5–7).

CircRNA derive from transcripts that are back-spliced and joined head-to-tail at the splice sites (6, 8). This covalent circularization of single stranded RNA molecules results in a novel backward fusion of two gene segments that can be of intronic and/or exonic origin (8). The formation of circRNA relies on complementary sequences in flanking introns that bring two splicing sites in close vicinity, and thus facilitate the back-splicing event (9). This circularization renders circRNA much more stable than linear RNAs (10). circRNA do not contain poly-A tails. As a consequence, they are not detected by the most widely used RNAseq methods that are based on poly-A selection. Our current knowledge of circRNA expression is therefore still at its infancy.

Several functions have been attributed to circRNA (11). They can serve as miRNA sponges (12, 13), or as transcriptional activators (14, 15). Furthermore, circRNA have been shown to segregate RNA binding proteins (16, 17), and can even become translated into proteins through cap-independent translation initiation (5, 18). CircRNA may also regulate the differentiation of HSCs (19). Indeed, circRNA expression has been described in several blood cells (7, 20–22). Together with recent reports on circRNA in neuronal and myocyte differentiation cells and in extracellular vesicles (5, 23), these findings prompted us to interrogate which circRNA are expressed in hematopoietic cells and whether the expression of circRNA alters during hematopoietic differentiation.

circRNA can be identified by their unique back-spliced junction, which results in chimeric reads alignment in the RNA-seq data. This feature distinguishes them from linear RNA (6); Figure 1A). Here, we used previously published transcriptome deep-sequencing data on primary human hematopoietic cells to define the expression pattern of circRNA during differentiation. This comprehensive analysis identified >59.000 circRNA in hematopoietic cells, including >14.000 newly annotated circRNA. We found that circRNA expression is cell-type specific and alters during differentiation. Furthermore, differentiated cells contain substantially higher levels of circRNA. We conclude that circRNA expression is widespread in hematopoietic cells, which warrants their further functional characterization.

**Figure 1:**
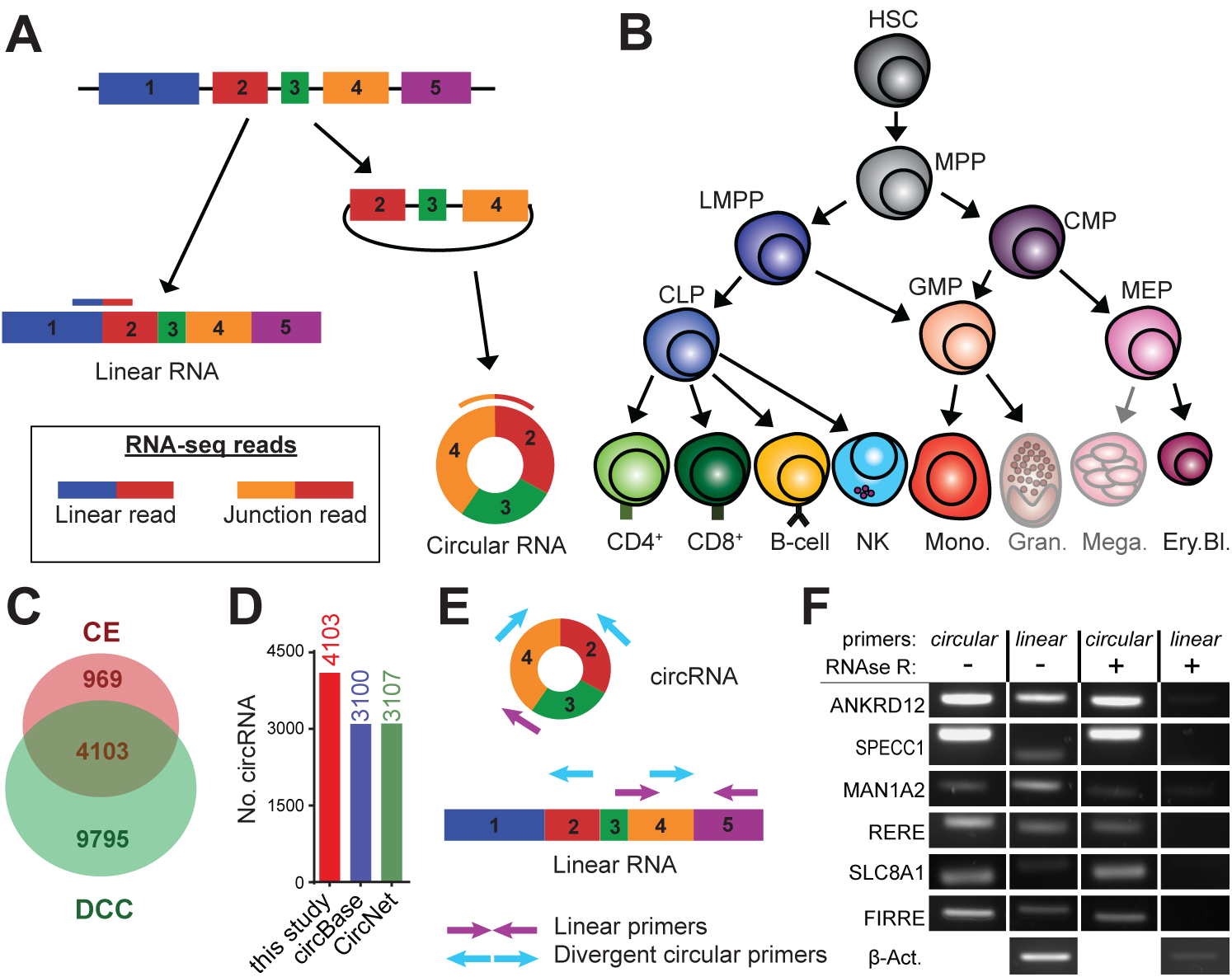
Circular RNA expression in hematopoietic cells. (**A**) Diagram presenting the forward-splicing of a linear RNA and the back-splicing of a circular RNA. (**B**) Diagram of hematopoietic differentiation (inspired by ref (24)). HSC: Hematopoietic stem cell, MPP: Multipotent progenitor, LMPP: lymphoid-primed multipotent progenitor, CLP: common lymphoid progenitor, CMP: common myeloid progenitor, GMP: Granulocyte-macrophage progenitor, MEP: Megakaryocyte-erythrocyte progenitor, CD4: CD4+ T cells, CD8: CD8+ T cells, NK: Natural killer cells, Mono.: Monocytes, Gran.: Granulocytes, Mega.: Megakaryocytes, Ery.Bl.: Erythroblasts. (**C**) The number of circRNA (≥ 2 junction reads in at least one sample of a specific cell population) detected in hematopoietic cells using DCC and CircExplorer2 (CE). (**D**) Numbers of circRNA detected by DCC and CE combined, compared to circRNA annotated in circBase and in circNet. (**E**) Scheme of primer design for detection by PCR of linear and circular RNA, respectively. (**F**) RNA extracted from human PBMCs was treated with RNAseR, or left untreated (control) prior to reverse transcription. β-actin mRNA primers were used as a linear transcript control. Data are representative of 4 donors.

## MATERIAL AND METHODS

### Data set selection

The SRA repository from NCBI was searched for datasets of human hematopoietic cells that contained paired-end data with long reads (>100nt), at least 20 million reads/sample, and that did not use Oligo-dT for cDNA preparation, or 5’ or 3’ capture methods for library preparation. The selected data set (accession GSE74246/SRP065216) from (24) used total RNA and the TruSeq/NEB Next Ultra library preparation kit. For granulocytes, RBC and platelets, we used datasets that used ribosomal RNA-depleted total RNA for sample preparation. Accession numbers for platelets: ERR335311, ERR335312 and ERR335313 (25); for RBCs: SRR2124299, SRR2124300, SRR2124301 and SRR2038798 (26, 27), and for granulocytes: ERR789064, ERR789082, ERR789195 and ERR789201 (28).

### circRNA identification and analysis

Data sets were aligned to the human genome (GRCh37 / hg19-release75) using STAR version 2.5.2b allowing for chimeric detection (*see Supplementary Methods*). The output file of STAR was analysed with DCC 0.4.4 (29) and CircExplorer2 (CE) 2.01 (30) to detect, filter and annotate circRNA (see supplementary method). DCC was used for detection of linear reads at the circRNA coordinates (using the option –G). For both CE and DCC, we used GRCh37/hg19 genome annotations.

circRNA expression was considered low confidence detection when at least 2 junction reads were found in at least 1 sample by both DCC and CE, and high confidence detection when at least 2 junction reads were found in all replicates of one specific cell type. Junction read counts were normalized to reads per million mapped reads (RPM). The maximal circRNA length was calculated with the exon length information provided by the CE annotations. These annotations were then used to calculate the first and last circularized exon.

For differential expression analysis of CircRNA in hematopoiesis cell, we calculated the log2 fold-change for each comparison of populations (see Supplemental method; (5, 23)). Hits were defined by log2FoldChange > 4 for high confidence circRNA. Kmeans clustering was performed with pheatmap 1.0.8 (31) with option *kmeans*_*k.* The number of 15 clusters was chosen after manual inspection of the circRNA clusters.

Circular over linear ratios (CLR) were calculated for the coordinates of high confidence circRNA (n=489) with the linear counts obtained with DCC (see Supplemental method). Linear counts were corrected for sequencing depth to obtain linear RPM. CLR was calculated by using circular RPM/linear RPM. Data analysis was performed in R (3.4.1) and R-Studio (1.0.143).

#### Plots and graphs

Heatmaps were generated in R using pheatmap 1.0.8. Plots and graphs were generated with ggplot2 (32) in R, or Graphpad PRISM version 7.0. Venn diagrams were produced with *http://bioinformatics.psb.ugent.be/beg/tools/venn-diagrams* from the University of Gent.

#### Cell population isolation

Peripheral blood mononuclear cells (PBMCs) were obtained according to the Declaration of Helsinki (seventh revision, 2013) from healthy volunteers with written informed consent (Sanquin, Amsterdam, the Netherlands). PBMCs were isolated by Lymphoprep density gradient separation (Stemcell Technologies). Specific cell types were isolated with CD4+ and CD8+ MACS beads for T cells, CD14+ beads for monocytes, CD56+ beads for NK cells, and CD34+ beads for CD34+ progenitors (purity > 98%, as determined by flow cytometry) according to the manufacturer’s protocols (Miltenyi). B cells were isolated with CD19+ Dynabeads and detach-a-beads (Invitrogen). Red blood cell fractionation was performed with a Percoll-Urografin gradient as previously described (33).

#### Flow cytometry

Red blood cell fractions were washed in PBS+1% bovine serum albumin (BSA, Sigma-Aldrich), and incubated for 30min at 4°C with anti-CD71-APC (Miltenyi; Clone AC102) and Thiazole orange (TO) (Sigma-Aldrich, 100 ng/mL). Cells were washed once with PBS+1% BSA and acquired on LSR Fortessa (BD). Data analysis was performed with FlowJo version X (Tree Star).

#### RNA extraction

*RNA was extracted* with Trizol (Life Technologies) according to the manufacturer’s protocol. RNA was resuspended in RNAse-free water. Half of the isolated RNA was treated for 15 min at 37°C with 8 units/μg RNAseR (Epicentre) in water supplemented with the digestion buffer provided by the manufacturer. RNAseR-treated and untreated control samples were purified with mini Quick-spin columns (Roche). cDNA preparation was performed with random hexamers using Super Script III reverse transcription (Invitrogen) according to the manufacturer’s protocol.

#### Primers and PCR

Primers were designed with CircInteractome (16) and with the NCBI primer blast tool (34), and manufactured by Sigma. Alternatively, the first and last 100nt of the circRNA junction were joined and used to blast for primer pairs. Primer sequences used to detect linear and circular RNA are found in *Supplementary Table 5.* PCRs were performed with ReddyMix PCR Master-Mix (Thermo-Fischer), and products were run on 2% agarose gels.

## RESULTS

### circRNAs are broadly expressed in hematopoietic cells

To determine which circRNA are expressed in hematopoietic cells, we used the RNA-seq data set of Corces et al. (24) that contained a broad range of hematopoietic progenitors and differentiated immune cells (Figure 1B). This data set was generated without selection for polyA+ mRNA, which allowed the identification and quantification of both circular and linear RNAs. Samples were included in the analysis when they reached a sequencing depth of at least 20M reads and when paired-end reads had a length of at least 100 bases. This cut-off resulted in 2-3 replicates per cell type.

We used two different tools for circRNA detection, i.e. DCC (29) and CircExplorer2 (CE) (30). Both tools allow for fast and accurate detection of circRNA by detecting back-splice junction reads of circRNA from RNA-seq data with a low false positive rate (reviewed in (35)). We used a cut-off for circRNA detection when at least 2 junction reads were measured in at least 1 sample of one specific population (7) (Figure 1C, Supplementary Table S1). This analysis identified 13.898 different circRNA with DCC and 5072 circRNA with CE. 4103 circRNA were detected with both tools, of which 3100 and 3107 were annotated in circBase (36) and CircNet (37) respectively (Figure 1D). 428 circRNA (10.4%) were newly identified circRNA that were not annotated in either data base (figure 1D). Only 16 circRNA (0.4%) of the 4103 circRNA derived from intronic junctions (Supplementary Figure S1A). The vast majority of the 4103 circRNA (4087 circRNA; 99.6%) contained exon-exon junctions.

We validated the expression of several highly expressed circRNA and the linear mRNA variants of the same gene by RT-PCR (Figure 1E) in human peripheral blood mononuclear cells (PBMC). Similar to the linear RNA products, all circRNA products identified with DCC or CE that we tested by RT-PCR were also detected in PBMCs (Figure 1F). The sensitivity of linear RNA, but not of circRNA, to the exonuclease ribonuclease R (RNAse R) confirmed the identity of the tested circRNA.

### circRNA generation, size and distribution in hematopoietic cells

We next determined which genes and which exons were used for the generation of circRNA. The distribution of the identified circRNA per chromosome was similar to the distribution of all coding and non-coding genes (Figure 2A). When analyzing the biological processes of circRNA-generating genes, they were enriched in housekeeping functions (Supplementary Figure S1B). The predicted numbers of exons used by circRNA is on average 5.96 exons as determined by CE based on the known linear exon usage, with a minimum of 1 exon and a maximum of 56 exons (Figure 2B). The calculated length of the circRNA based was on average 1057 nucleotides (Figure 2C).

**Figure 2:**
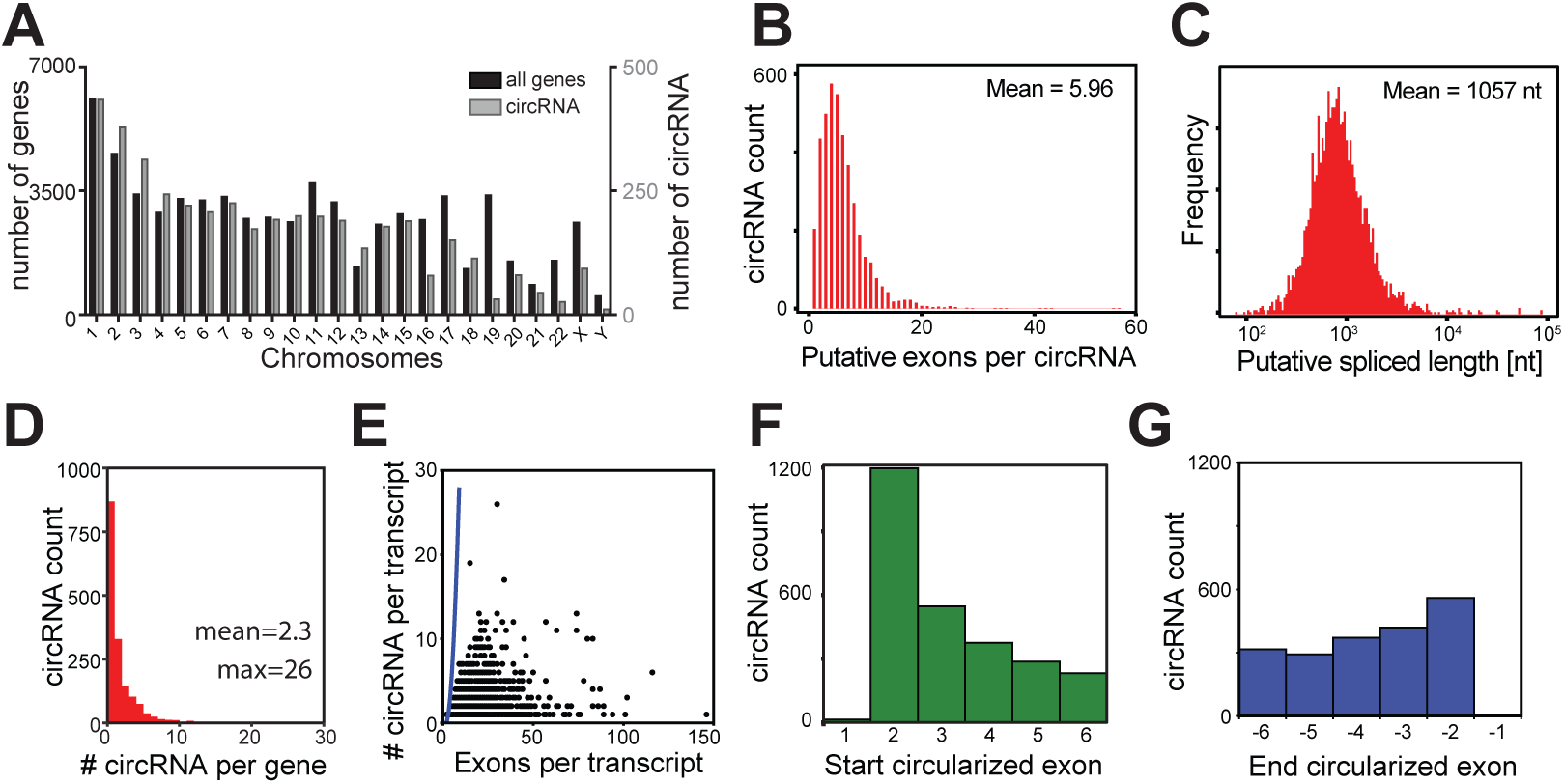
Characterization of circRNA detected in hematopoietic cells. (**A**) Distribution of circRNA (in grey) and known genes from hg19/GRCh37 (in black) per chromosome. (**B, C**) Putative exon usage (**B**) and maximal putative length (**C**) of the 4103 circRNA detected in hematopoiesis, as determined by the start and end position of circRNA and splicing annotations of the RNA linear transcript. (**D**) Number of detected circRNA per gene (n=4103). (**E**) Relation between the number of circRNA per transcript and the number of exons of that transcript. Blue line represents the calculated maximum circRNA number per transcript, if all exon combinations were used for circularization. (**F-G**) Distribution of the first (**F**) and the last (**G**) 6 exons used for circularization from the linear RNA in circRNA. 1197/4103 (29.2%) circRNA use the second exon as start exon, and 559/4103 (13.6%) use the one but last exon as end exon.

We next interrogated how many circRNA variants are generated from a gene. To this end, we calculated all different back-splice junction reads found per gene in hematopoietic cells. This analysis revealed that 930 genes (53%) generated only one circRNA variant. However, the overall expression of circRNA variants is between 1 and 26 variants per gene with a mean of 2.3 circRNA/gene (Figure 2D). If circRNA would be a random event, the maximum possible number of circRNA should increase near-exponentially as the number of exons per transcripts increases (Figure 2E, blue line). There was, however, no overt correlation between the detected number of circRNA variants per gene and the number of exons per transcripts, suggesting that the generation of circRNA is not a random event (Figure 2E; compare the data points with blue line).

Finally, we questioned which exon of the linear transcript is used by circRNA as the starting exon for circularization. In line with the canonical splicing rules (38), circRNA in hematopoietic cells barely use the first exon (Figure 2F). Rather, circRNA favor the second exon of a transcript as starting exon, which is found in 1197/4103 circRNA (29.2%) (Figure 2F). We also determined which exon is used for back-splicing, referred to as the end circularized exon. Again, the last exon of a linear transcript was not used (Figure 2G). However, we could not detect a clear preference for a specific end exon (Figure 2G).

### CircRNA expression in hematopoiesis alters upon differentiation

We next interrogated whether the expression of circRNA alters during hematopoietic differentiation. We focused on circRNA that were identified with high confidence by both DCC and CE with at least 2 junction reads (Figure 1C), and that were found in each biological replicate of one specific cell population. This analysis yielded 489 high confidence circRNA (Figure 3A, Supplementary Table S1).

**Figure 3:**
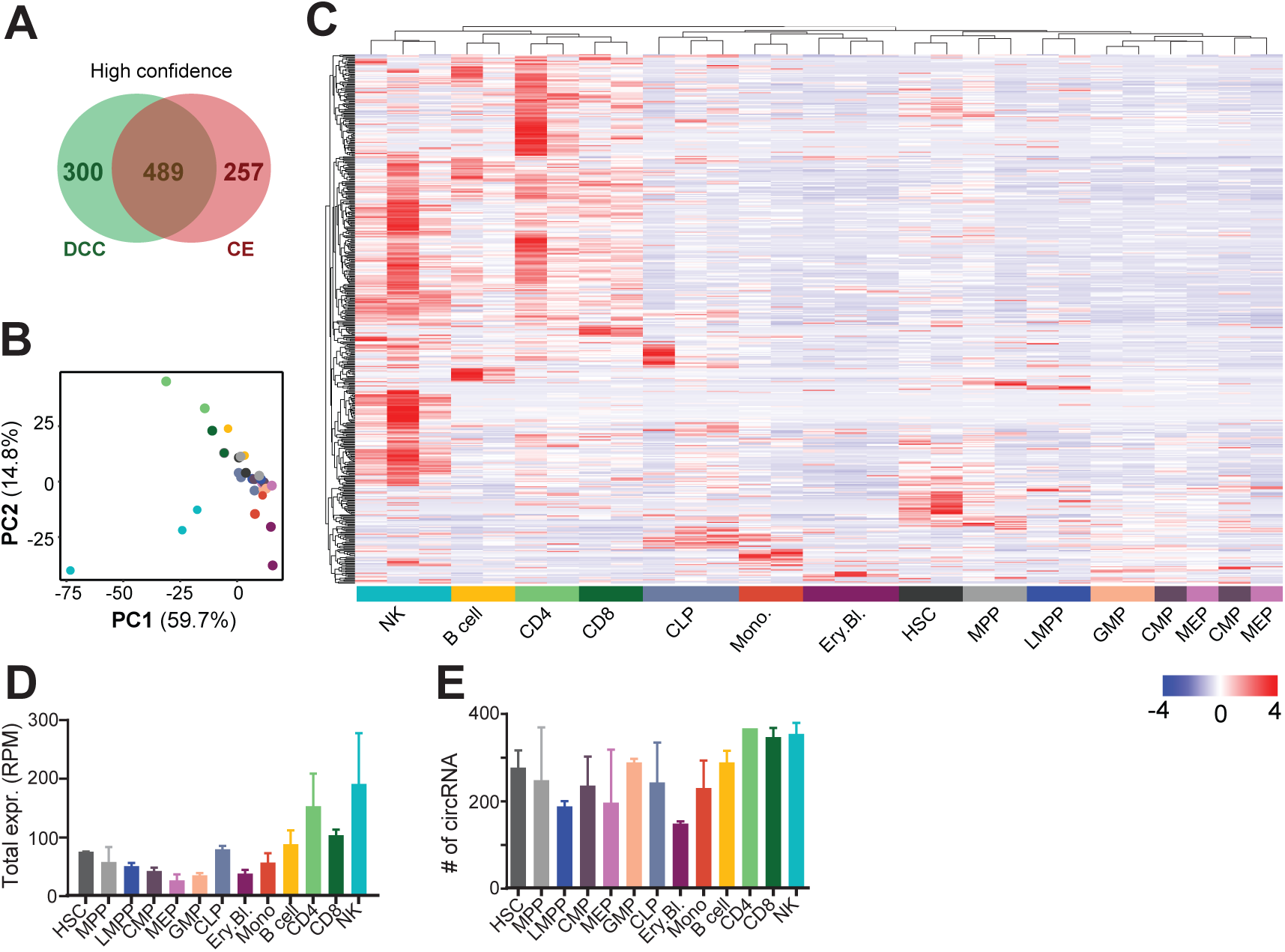
circRNA expression in hematopoietic cells. (**A**) Number of high confidence circRNA detection, as defined by ≥ 2 junction reads in each replicate of one specific cell population). 33 out of the 489 circRNA were newly identified. (**B-C**) PCA plot (**B**) and (**C**) unsupervised Pearson correlation clustering (row-scaled RPM) of the circRNA expression in the hematopoietic populations depicted in a heat map (n=2-3 per cell type). (**D-E**) Sum of different circRNA expressed per cell type (**D**), and of the overall number of circRNA expressed per population (**E**). n=2-3, error bar depicts standard deviation. RPM: reads per million mapped reads.

Principal component analysis (PCA) of the circRNA expression accurately recapitulated the previously described distribution of the linear mRNA in hematopoietic cell populations (24). HSC and progenitor cells are located at the intersection of the differentiated myeloid and lymphoid cells (Figure 3B). Differentiated myeloid cells clustered with each other, away from the progenitors, and in opposite directions as the mature lymphoid cells (Figure 3B). Unsupervised clustering based on circRNA expression also showed a clear clustering of HSCs with progenitors in a heat map, away from differentiated cell populations (Figure 3C). We observed that lymphocytes had the highest levels of circRNA expressed, based on the sum of all circRNA RPM (Figure 3D). The number of distinct circRNA, however, was similar in various cell populations, indicating that the abundance of circRNA, but not the variety of circRNA, was increased in B cells, CD4^+^ and CD8^+^ T cells, and in NK cells (Figure 3D,E).

### CircRNA expression during hematopoietic differentiation is cell-type specific

To determine the cell-type specific expression of circRNA, we looked in more detail at circRNA of the highly confident 489 circRNA detected in Figure 3, which displayed a log2FoldChange ≥4 (≥16 times enriched) in any comparison between two populations. This cut-off identified 187 circRNA (Figure 4A, Supplementary Table S2). K-means clustering (*k*=15) showed cell-type specific circRNA expression pattern (Figure 4A, B).

**Figure 4:**
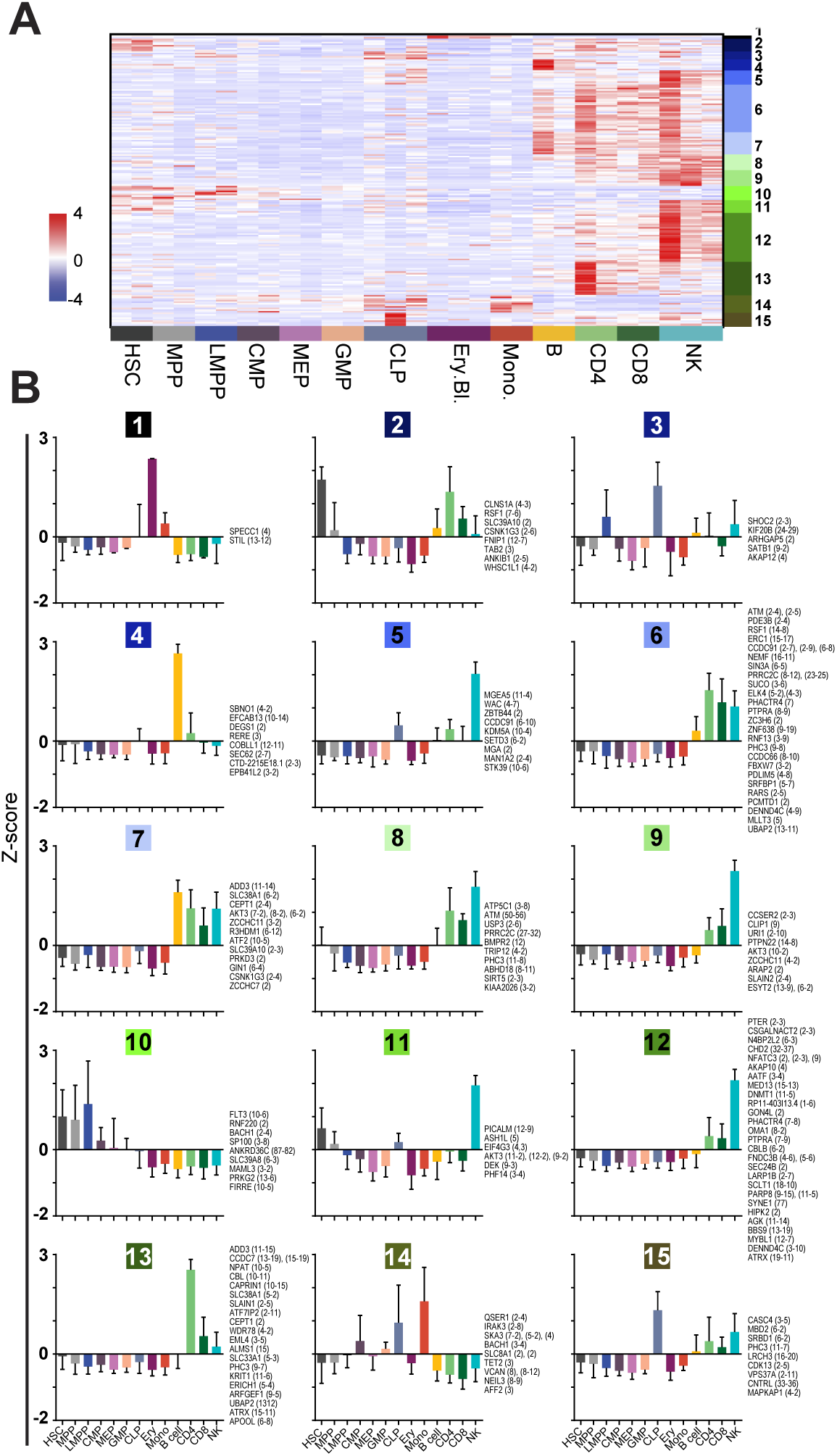
circRNA expression is cell-type specific. (**A**) Heatmap of circRNA that are differentially expressed, with a cut-off of log2FoldChange ≥4 (≥16 times enriched) in at least one cell population. At the right, the 15 K-means clusters are depicted (n=187, row-scaled RPM). (**B**) K-means clusters (*k*=15) depicting the relative expression (Z-score) across the hematopoietic cell populations. circRNA gene names with the exon usage in brackets are depicted for each cluster on the right of the graph.

Clusters 2, 3, 10, 11, and 15 include circRNA expressed by the progenitors (Figure 4B). CircRNA in cluster 10 were mainly expressed in early progenitors, i.e. HSCs, MMPs, and LMPPs. Cluster 2 circRNA were more restricted to HSCs and MPPs, but also showed circRNA expression in lymphoid cells. Clusters 3 and 15 showed the signature of CLPs, concomitant with low expression in differentiated lymphoid cells. Cluster 11 circRNA were shared between HSC and MPPs, and with NK cells.

Lymphoid cells specific circRNA were represented by clusters 6-9 (Figure 4B). Cluster 4 was specific for B cells, cluster 13 for CD4^+^ T cells, and cluster 5 for NK cells. Cluster 9 and 12 showed a mixed lymphoid signature that was enriched for NK cells. Myeloid cell specific circRNA were present in cluster 1 and 14 (Figure 4B). Cluster 1 was erythroid specific and 14 mostly monocyte-specific with some background expression in CMP and CLP. We validated the distribution of several circRNA in blood cell subsets. Indeed, circ-FNDC3B (exon 5-6) was broadly expressed with the highest expression levels in NK cells. Circ-ELK4 (exon 4-3) and circ-MYBL1 (exon 12-7), and the newly identified circ-SLFN12L (cluster 2-3) show the highest expression in T cells and NK cells (Supplementary Figure S1C).

Interestingly, different circRNA variants generated from the same gene were detected in different clusters (Figure 4B). For example, circ-BACH1 (exon 3 to 4) was preferentially expressed in monocytes (cluster 14), whereas circ-BACH1 (exon 2 to 4) was increased in HSCs and MPPs (cluster 10). This finding was confirmed by RT-PCR (Supplementary Figure S1D). Likewise, different variants of circ-AKT3 (clusters 7, 9, and 11) and circ-CCDC91 circRNA (clusters 5 and 6) were expressed across lymphoid cells. In conclusion, our analysis shows that the expression of circRNA is cell-type specific.

### The ratio of circRNA over linear RNA alters with differentiation

We next investigated how the expression levels of circRNA compares with that of the corresponding linear RNA. Because circRNA detection is restricted to the junction reads, the levels of linear mRNA expression are also defined by comparing the number of reads at the start and at the end position of the circRNA (Figure 5A). DCC analysis corrects for the double counting of linear reads (start + end; (29)), and allows us to calculate the circular-over-linear ratio (CLR) on the 489 circRNA expressed in hematopoietic cells (Figure 3, 5A).

**Figure 5:**
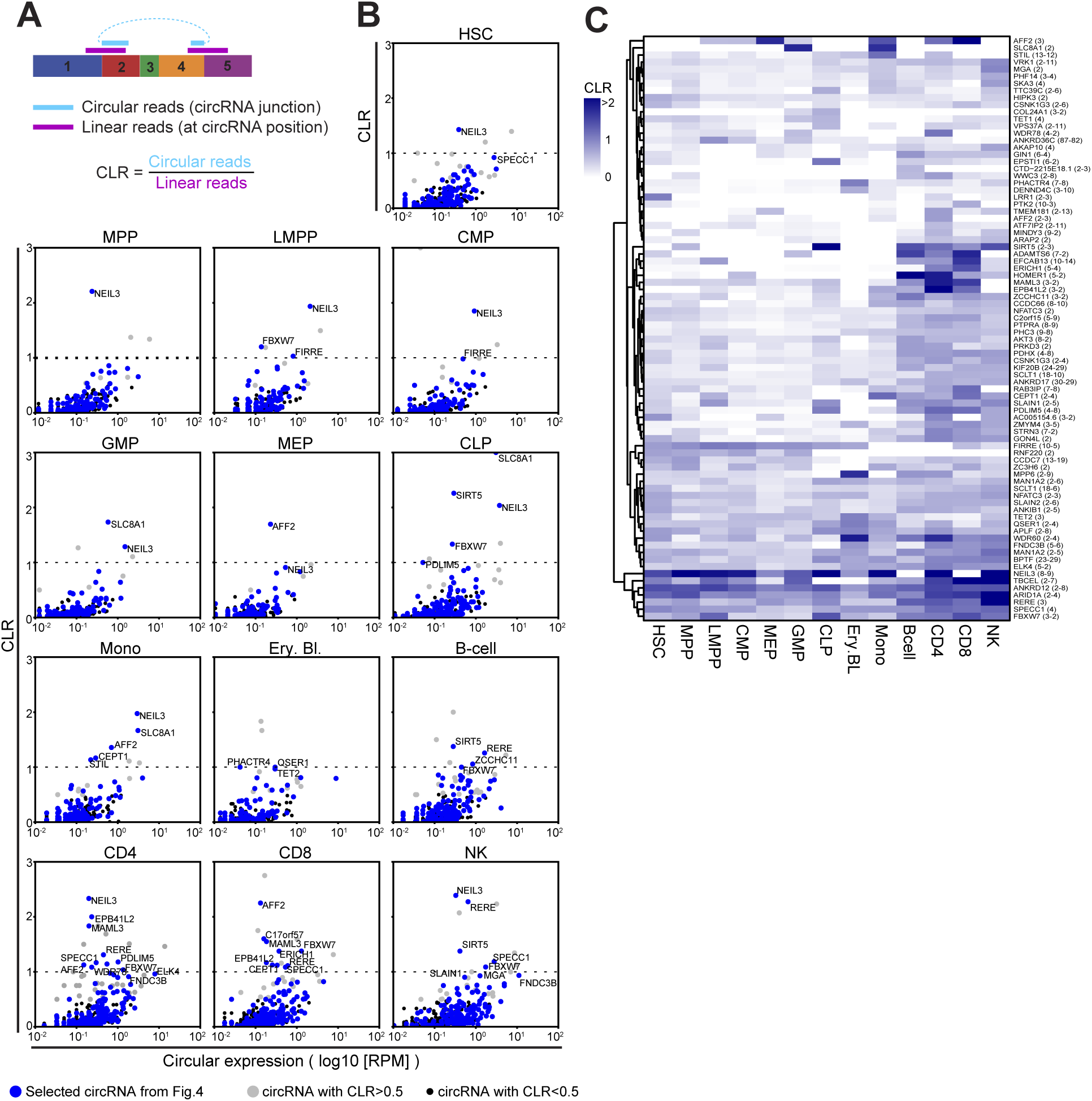
CircRNA usage alters during hematopoietic differentiation. (**A**) Scheme showing how the circular-to-linear ratio (CLR) is computed using DCC (29). (**B**) CLR of circRNA from Figure 3 for each population against the number of reads of the circRNA (log10 (RPM)). CLR>1 indicates that the sample contains more transcripts for the circRNA than the corresponding linear transcript. Blue dots are the 187 differentially expressed circRNA from Figure 4, grey dots are circRNA with a CLR>1, and black dots are circRNA with CLR<1. (**C**) Heatmap showing the average CLR within a population, of circRNA having CLR>0.5 in at least one of the hematopoietic populations.

We plotted the CLR against the circRNA expression for each hematopoietic population. 82 circRNA showed a CLR>0.5, indicating that these circRNA reached at least 50% of the levels of the linear RNA (Figure 5B, gray dots). The 187 circRNA used to determine cell-type specific circRNA expression in Figure 4 followed the same expression pattern (Figure 5B, blue dots). The expression level for several circRNA reached even higher expression levels than the corresponding linear RNAs (CLR >1; Figure 5B, Supplementary Table S3). The number of circRNA with a CLR>1 was highest in differentiated lymphoid cells, i.e. B cells, T cells and NK cells.

We next determined if the ratio between circular and linear RNAs alters during hematopoietic differentiation. To this end, we plotted all circRNA with a CLR >0.5 (n=82) in a heat map (Figure 5C). Some circRNA like circ-NEIL3 (exon 8-9) or circ-ANKRD12 (exon 2-8) retained a high CLR in all cell populations. In contrast, the CLR of circ-SLC8A1 (exon 2) was only high in GMPs and in monocytes. The CLR of circ-AFF2 (exon 3) was high in MEPs and monocytes as well as in T cells (Figure 5C). Circ-RERE (exon 3) showed the highest CLR in NK cells, but was also found in B cells and T cells, albeit with lower CLRs (Figure 5C). Combined, these findings show that the use of circRNA and their correlation to the linear RNA is cell-type specific and changes during hematopoiesis.

### Red blood cells and platelets have high expression levels of circRNA

Maturation of red blood cells is accompanied with an increase in heterochromatin and condensation of the nucleus, followed by extrusion of the nucleus. Platelets are shed from megakaryocytes. Both red blood cells and platelets undergo their final steps of maturation in absence of active gene transcription (39). To investigate the expression levels of circRNA in RBC and platelets, we first used publicly available RNA expression data of platelets, red blood cells (RBC), and granulocytes isolated from peripheral blood (25–28). By combining DCC and CE and using the low-confidence cut-off of 2 junction reads in at least 1 replicate of one specific cell population, we identified 59.011 circRNA, of which 28.841 and 42.082 were annotated in circBase and CircNet, respectively (Figure 6A). Over 14.100 circRNA (23.9%) were thus newly identified in this data set. Of the 59.011 identified circRNA, platelets express the highest numbers of circRNA (47.654), followed by RBCs (27.409) and granulocytes (8.925) (Supplementary Table S4). With the high confidence cut-off of 2 junction reads in each biological replicate of one cell type, we detected 10.729 circRNA in platelets, 5.878 circRNA in RBCs, and 1.989 circRNA in granulocytes (Figure 6B, Supplementary Table S4). Of these, both shared and cell-type specific circRNA were detected (Figure 6B). Platelets not only contained the highest numbers of different circRNA, but also clearly outnumbered RBCs and granulocytes with the overall RPM of circRNA RPM (Figure 6C).

**Figure 6:**
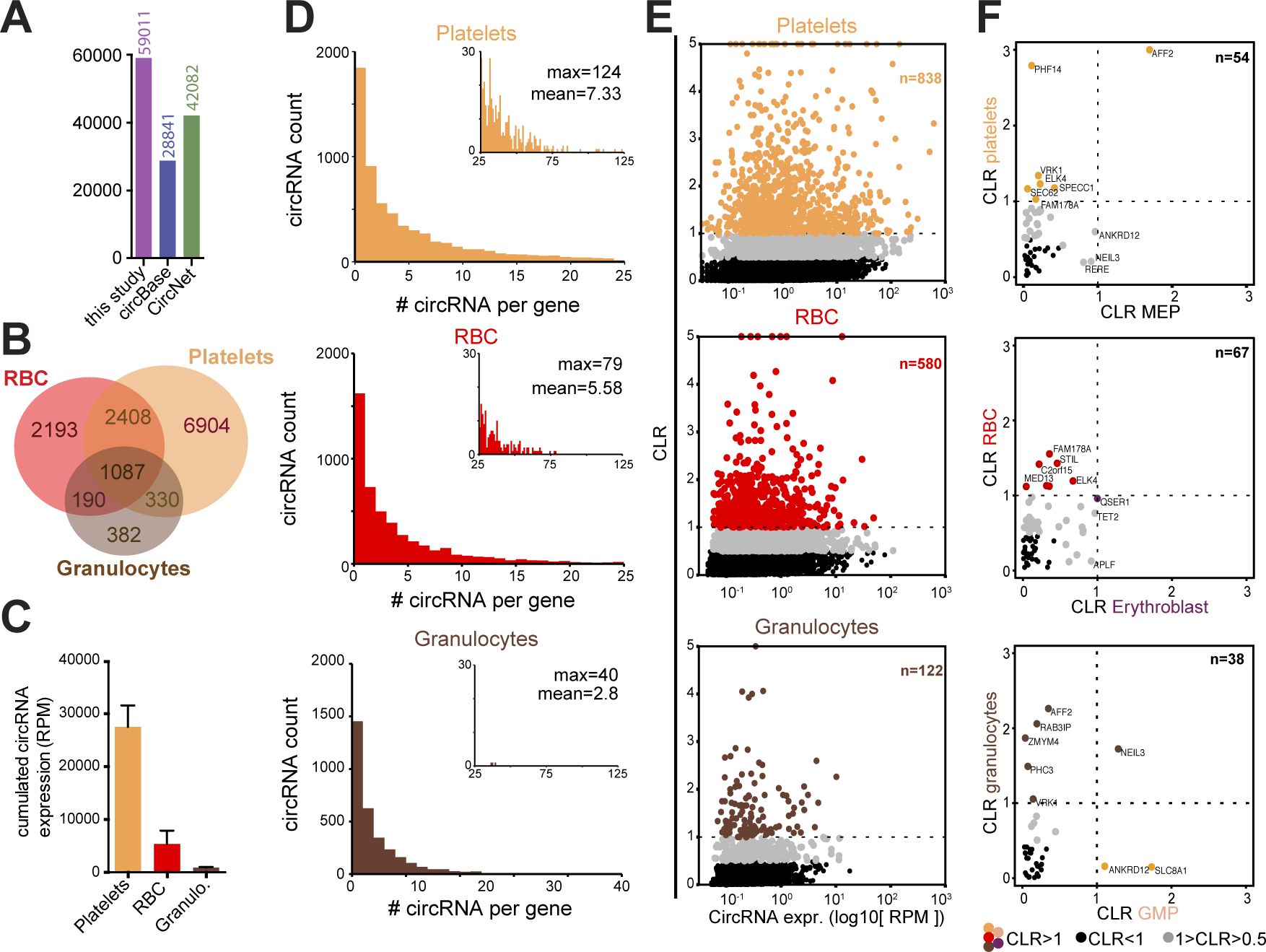
Red blood cells and platelets express high levels of circRNA. (**A**) CircRNA detection (low confidence; i.e. ≥ 2 junction reads in at least one sample) in granulocytes, red blood cells (RBC) and platelets, compared to annotations in circBase and CircNet databases. (**B**) circRNA expression with high confidence (≥ 2 junction reads in all replicate of one population). (**C**) The sum of circRNA detected with high confidence per cell type (RPM). (**D**) Number of circRNA detected per gene in platelets (top panel), RBC (middle panel) and granulocytes (lower panel). The insert histograms show circRNA found with more than 25 circRNA/gene. Numbers depict Maximum circRNA per gene (max) and the average circRNA per gene (mean). (**E**) Circular-to-linear ratio (CLR) plotted against the circRNA expression (log10 (RPM)). Black dots show circRNA with CLR<1 and colored dots show circRNA with CLR>1, for platelets (top panel), RBC (middle panel) and granulocytes (lower panel). (**F**) CLR of circRNA detected in both progenitor cells and differentiated cells for platelets with MEP (n=169 circRNA, top panel), RBC with erythroblasts (n=177 circRNA, middle panel) and granulocytes with GMP (n=104 circRNA, lower panel).

We next determined the number of circRNA variants that are expressed per gene in the three different cell types (Figure 6D). Of note, none of the three cell types were included in the data set of Figure 1-5. Granulocytes expressed on average 2.8 circRNA per gene with a maximum of 40 circRNA/gene, which resembled the numbers of circRNA measured per gene in hematopoietic cells (Figure 2D). Conversely, RBCs and platelets express on average 5.58 and 7.33 circRNA per gene, respectively, with a maximum of 79 for RBCs and 124 for platelets (Figure 6D). Thus, the relative expression levels of circRNA and the number of variants expressed per gene in enucleated RBCs and platelets exceed those of all other hematopoietic cell types.

The high prevalence of circRNA in platelets and RBCs is also reflected by the high CLR (Figure 6E). Concomitant with the loss of *de novo* transcription in enucleated cells, RBCs and platelets contain high levels of circRNA, and a significant number of circRNA have a high CLR. 7.8% of the circRNA expressed in platelets had a CLR > 1 (n=838). Similarly, RBCs had 9.87% circRNA with a CLR >1 (n=580), and 6.13% (n=122) in granulocytes (Figure 6E).

We then determined how the circRNA expression pattern of these three cell types corresponded to the circRNA usage in the respective progenitor. The hematopoiesis dataset (24) and the differentiated myeloid cells (25–28) were two different data sets generated with different methods, i.e. total RNA and with ribosomal RNA depletion, respectively. We therefore used the CLR of each circRNA for comparison of progenitors and differentiated cells, i.e. MEP and platelets; erythroblasts and RBCs; and GMP and granulocytes. We included the circRNA and the corresponding linear mRNAs in this analysis that had at least 0.1 RPM each (Figure 6F; Supplementary Table S4). The circRNA that have a CLR>0.5 (gray) or of a CLR>1 (colored) are more prevalent in mature platelets, RBCs and granulocytes than in the precursor populations (Figure 5F). In addition, the circRNA with a CLR>0.5 in mature cells barely overlap with those of the precursor cells (Figure 5F). Combined, we conclude that circRNA are highly expressed in differentiated myeloid cells, and that the highest relative levels are detected in the enucleated RBCs and platelets.

### CircRNA expression retains in aging RBCs

RBCs have a life span of 120 days, and during this period RBCs age (40). We therefore analysed if and how the linear and circular RNA expression altered during RBC maturation and ageing. To separate reticulocytes and young erythrocytes from old erythrocytes we used a Percoll-Urografin gradient ((33), Figure 7A). The maturation status of reticulocytes and the erythrocytes was confirmed with CD71 expression and the nucleic acid content with Thiazole Orange (TO) by flow cytometry (Figure 7A). Upon ageing, the expression of CD71 is lost (Figure 7B) and the TO staining (Figure 7C) is decreased.

**Figure 7:**
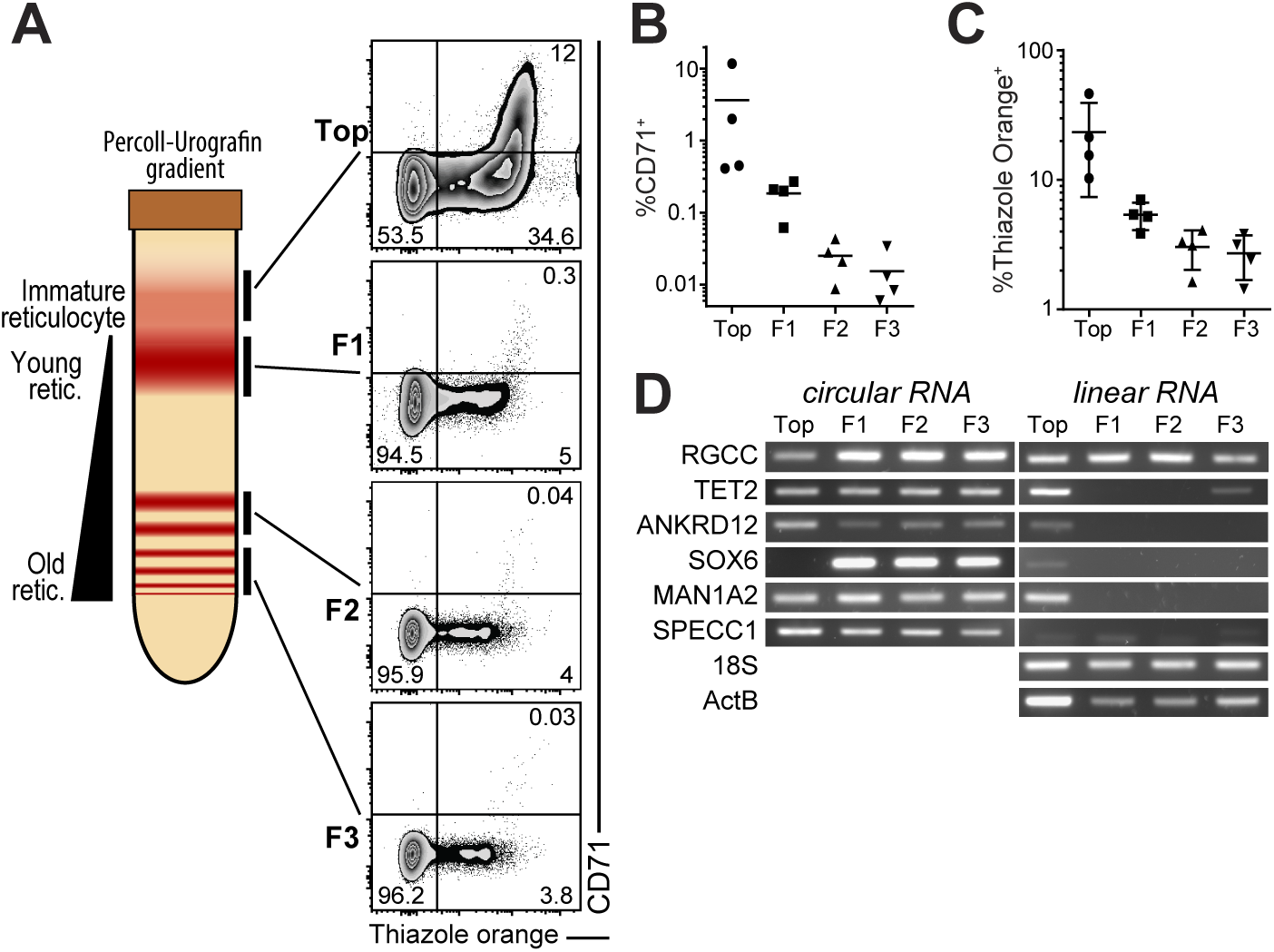
circRNA are conserved throughout red blood cell aging. (**A**) Left panel shows a representation of a Percoll-Urografin gradient separation after centrifugation. The immature reticulocyte fraction is found in the top fraction (bellow a neutrophil-rich ring), the lower fractions (F1, F2, F3) separate the aging red-blood cells from young to old (top to bottom). Right panel shows a flow-cytometry measurement of the fractions for CD71 expression and nucleic acid (thiazole orange) staining (representative of 4 donors). (**B-C**) Graph showing the percentage of CD71+ (**B**) and thiazole orange + (**C**) in the fractions from (A) (n=4). (**D**) RNA extracted from the fractions in (A) were reverse transcribed and circRNA and linear RNA detection was performed using PCR. 18S and Beta-actin (ActB) were used as a linear control. (D) is representative of 3 donors.

We used these four fractions of red blood cells to detect RBC-specific circRNA with relatively high expression and CLR over 0.5 and their linear mRNAs by RT-PCR (Figure 7D). During ageing, only the highly stable *β*-*ACTIN mRNA*, and *RGCC* mRNA is detected in mature erythrocytes (Figure 7D, right panel). All other measured linear mRNAs lose their expression upon differentiation (Figure 7D, right panel). In contrast, circ-TET2 (exon 3), circ-ANKRD12 (exon 2-8), circ-MAN1A2 (exon 2-5), and SPECC1 (exon 4) remained stably detectable in young, and in old erythrocytes (Figure 7B, left panel). Interestingly, whereas *SOX6* mRNA was only detectable in young reticulocytes, circ-SOX6 (exon 8-9) was specifically detected in mature reticulocytes (Figure 7B). Thus, circRNA expression is stably maintained during RBC ageing, suggesting that circRNA may be involved in RBC function.

In conclusion, we present here an analysis of circRNA in hematopoietic cells. We demonstrate, that circRNA expression is cell type specific, and that mature cells both of lymphoid and of myeloid origin contain more circRNA.

## DISCUSSION

CircRNA can drive a subset of cellular functions (5, 12, 41), yet their role in gene regulation and expression in human hematopoietic cells is not well understood. To our knowledge, we here provide the first comprehensive analysis of circRNA expression during hematopoiesis. We identified over 14.000 novel circRNA, which significantly extends the list of currently annotated circRNA in the human transcriptome. How specific these newly identified circRNA are for hematopoietic cells is yet to be determined. Because the identified circRNA are formed both from genes with general functions and from hematopoietic specific genes, and because the CLR indicates a cell-type specific enrichment of circRNA, we postulate that at least some of the identified circRNA should be cell-type specific.

Similar to differentiating neurons and myocytes (5, 23) the expression levels of circRNA increase during hematopoietic differentiation. This increase could be a mere result of accumulation of stable circRNA during differentiation. However, we show here that some circRNA are preferentially expressed in progenitors. For example, the expression levels of circ-FIRRE (exon 5-10) was high in all progenitors except from CLPs, and it was significantly reduced upon differentiation (Figure 4B). Another product of the *FIRRE* gene, the lncRNA FIRRE was shown to regulate part of the pluripotency of embryonic stem cells (ESC) via Repeat RNA Domains (RRD). Intriguingly, we observed that circ-FIRRE (5–10) also contains all RRDs described for the LncRNA FIRRE. It is therefore tempting to speculate that some of the effects that are attributed to the lncRNA FIRRE could also be exerted by the circular FIRRE RNA.

Interestingly, the exon usage for circRNA generation - as we describe for circ-BACH1 - can alter during differentiation. Several studies showed that alternative splicing during hematopoietic differentiation is primarily driven by exon skipping (2, 39). The original hypothesis was that skipped exons were degraded via non-sense mediated decay. However, these exons can also be used for the formation of circRNA (42). As exon skipping is increased during late erythropoiesis (39), and circRNA expression is then increased, it is tempting to speculate that exon skipping contributes to the production of circRNA in erythrocytes. However, exon skipping cannot be the sole and perhaps even not the major source of circRNA. Many circRNA contain several exons, and the generation thereof appears cell type specific and differentiation dependent. In addition, a subset of circRNA display an increased expression over the linear variant. Combined, these data suggest a highly orchestrated production of circRNA. Future investigations should reveal how the formation of circRNA relates to the exon skipping, or to other regulatory events.

Platelets and erythrocytes contain high levels of circRNA. This can at least in part be explained by the degradation of linear transcripts (26). Interestingly, the circRNA expression of erythroblast and MEP does not fully overlap with the differentiated RBCs and platelets. This again suggests that in addition to accumulation of stable circRNA in enucleated cells also the generation of circRNA may alter during differentiation. Furthermore, platelets release specific circRNA in extracellular vesicles (43), which may alter the composition of the remaining circRNA. This specific release also suggests that circRNA may be involved in platelet-associated cellular processes.

The function of circRNA is to date not well understood (44). Whereas some circRNA can serve as a miRNA sponge (45), this does not apply to most of the currently known circRNA (13, 46). CircRNA may also serve as sponges for RNA-Binding Proteins (11, 17), or crosstalk with the transcriptional machinery (14, 15). Recent studies show that circRNA can be also translated into functional protein (5, 18, 47). Thus, the function of circRNA is most probably diverse. Here we provide a comprehensive analysis of circRNA in hematopoietic cells. Future studies will reveal how circRNA can contribute to the regulatory processes in hematopoietic cells, and possibly in the make up of the proteome. Furthermore, circRNA may be a useful tool to distinguish diseased from healthy hematopoietic cells and be used as potential biomarkers, as previously suggested for several other types of cancers (48).

## DATA AVAILABILITY

All bioinformatics tools used in this study are described in the supplementary method. The following datasets were used in this study: SRP065216, ERR335311, ERR335312, ERR335313, SRR2124299, SRR2124300, SRR2124301, SRR2038798, ERR789064, ERR789082, ERR789195, ERR789201, and are available on SRA (https://www.ncbi.nlm.nih.gov/sra).

## SUPPLEMENTARY DATA

**Supplemental Figure 1:** (**A**) Number of circRNA and circular intronic (ciRNA) from the low confidence circRNA in hematopoietic cells from CE annotations (≥ 2 junction reads in at least one sample of a specific cell population). (**B**) Gene Ontology (GO) analysis of the circRNA depicted in (**A**). (**C**) Detection of selected circRNA by RT-PCR from different from hematopoietic populations (each sample representative of 2 donors). (**D**) Expression two circRNA variants for BACH1 in different hematopoietic cells using different exon junctions, i.e. (2–4) and (3–4).

**Supplementary Table 1:** Full list of circRNA detected by DCC, CE and detected by both tools (see different tabs). Low confidence (n=4103), and high confidence (n=489) are included in separate tabs.

**Supplementary Table 2:** List and details of differentially expressed circRNA from Figure 4 (n=187).

**Supplementary Table 3:** List of circRNA and their CLR ratio from Figure 5B, the populations are found in different tabs.

**Supplementary Table 4:** circRNA detected in platelets, RBC and granulocytes by DCC and CE combined from Figure 6. Low confidence and high confidence circRNA with their CLR are in tabs. Last tabs are listing the circRNA with their CLR ratio detected in platelets vs MEP, RBC vs erythroblasts, granulocytes vs GMP from figure 6F.

**Supplementary Table 5:** PCR primer sequence used for detection of linear and circular RNA.

## ACKNOWLEDGMENTS

We would like to thank S. Heshusius and K. Moore for input on bioinformatics, E. Heideveld, M. Hansen, P-P. Unger, and R. Temming for help in isolating monocytes, platelets, B-cells, and NK cells, respectively, and the department of hematopoiesis for providing isolated CD34+ cells.

## REFERENCES

1. Orkin,S.H. and Zon,L.I. (2008) Hematopoiesis: An Evolving Paradigm for Stem Cell Biology. Cell, 132, 631–644.

2. Goode,D.K., Obier,N., Vijayabaskar,M.S., Lie-A-Ling,M., Lilly,A.J., Hannah,R., Lichtinger,M., Batta,K., Florkowska,M., Patel,R., et al. (2016) Dynamic Gene Regulatory Networks Drive Hematopoietic Specification and Differentiation. Dev. Cell, 36, 572–587.

3. Luo,M., Jeong,M., Sun,D., Park,H.J., Rodriguez,B.A.T., Xia,Z., Yang,L., Zhang,X., Sheng,K., Darlington, G.J., et al. (2015) Long non-coding RNAs control hematopoietic stem cell function. Cell Stem Cell, 16, 426–438.

4. Bissels,U., Bosio,A. and Wagner,W. (2012) MicroRNAs are shaping the hematopoietic landscape. Haematologica, 97, 160–167.

5. Legnini,I., Di Timoteo,G., Rossi,F., Morlando,M., Briganti,F., Sthandier,O., Fatica,A., Santini,T., Andronache,A., Wade,M., et al. (2017) Circ-ZNF609 Is a Circular RNA that Can Be Translated and Functions in Myogenesis. Mol. Cell, 66, 22–37.e9.

6. Memczak,S., Jens,M., Elefsinioti,A., Torti,F., Krueger,J., Rybak,A., Maier,L., Mackowiak,S.D., Gregersen,L.H., Munschauer,M., et al. (2013) Circular RNAs are a large class of animal RNAs with regulatory potency. Nature, 495, 333–338.

7. Maass,P.G., Gla,P., Memczak,S., Dittmar,G., Hollfinger,I., Schreyer,L., Sauer,A. V and Toka,O. (2017) A map of human circular RNAs in clinically relevant tissues. 10.1007/s00109-017-1582-9.

8. Chen,L.L. (2016) The biogenesis and emerging roles of circular RNAs. Nat. Rev. Mol. Cell Biol., 17, 205–211.

9. Ivanov,A., Memczak,S., Wyler,E., Torti,F., Porath,H.T., Orejuela,M.R., Piechotta,M., Levanon,E.Y., Landthaler,M., Dieterich,C., et al. (2015) Analysis of intron sequences reveals hallmarks of circular RNA biogenesis in animals. Cell Rep., 10, 170–177.

10. Jeck,W.R., Sorrentino,J.A., Wang,K., Slevin,M.K., Burd,C.E., Liu,J., Marzluff,W.F. and Sharpless,N.E. (2013) Circular RNAs are abundant, conserved, and associated with ALU repeats. Rna, 19, 141–157.

11. Hentze,M.W. and Preiss,T. (2013) Circular RNAs: Splicing’s enigma variations. EMBO J., 32, 923–925.

12. Piwecka,M., Glažar,P., Hernandez-Miranda,L.R., Memczak,S., Wolf,S.A., Rybak-Wolf,A., Filipchyk,A., Klironomos,F., Cerda Jara,C.A., Fenske,P., et al. (2017) Loss of a mammalian circular RNA locus causes miRNA deregulation and affects brain function. Science (80-.)., 8526, eaam8526.

13. Guo,J.U., Agarwal,V., Guo,H. and Bartel,D.P. (2014) Expanded identification and characterization of mammalian circular RNAs. Genome Biol, 15, 409.

14. Zhang,Y., Zhang,X.O., Chen,T., Xiang,J.F., Yin,Q.F., Xing,Y.H., Zhu,S., Yang,L. and Chen,L.L. (2013) Circular Intronic Long Noncoding RNAs. Mol. Cell, 51, 792–806.

15. Li,Z., Huang,C., Bao,C., Chen,L., Lin,M., Wang,X., Zhong,G., Yu,B., Hu,W., Dai,L., et al. (2015) Exon-intron circular RNAs regulate transcription in the nucleus. Nat. Struct. Mol. Biol., 22, 256–264.

16. Dudekula,D.B., Panda,A.C., Grammatikakis,I., De,S., Abdelmohsen,K. and Gorospe,M. (2016) CircInteractome: A web tool for exploring circular RNAs and their interacting proteins and microRNAs. RNA Biol., 13, 34–42.

17. Li,B., Zhang,X.-Q., Liu,S.-R., Liu,S., Sun,W.-J., Lin,Q., Luo,Y.-X., Zhou,K.-R., Zhang,C.-M., Tan,Y.-Y., et al. (2017) Discovering the Interactions between Circular RNAs and RNA-binding Proteins from CLIP-seq Data using circScan. bioRxiv.

18. Pamudurti,N.R., Bartok,O., Jens,M., Ashwal-Fluss,R., Stottmeister,C., Ruhe,L., Hanan,M., Wyler,E., Perez-Hernandez,D., Ramberger,E., et al. (2017) Translation of CircRNAs. Mol. Cell, 66, 9–21.e7.

19. Bonizzato,A., Gaffo,E., Te Kronnie,G. and Bortoluzzi,S. (2016) CircRNAs in hematopoiesis and hematological malignancies. Blood Cancer J., 6, e483–12.

20. Durek,P., Nordström,K., Gasparoni,G., Salhab,A., Kressler,C., de Almeida,M., Bassler,K., Ulas,T., Schmidt,F., Xiong,J., et al. (2016) Epigenomic Profiling of Human CD4+T Cells Supports a Linear Differentiation Model and Highlights Molecular Regulators of Memory Development. Immunity, 45, 1148–1161.

21. Memczak,S., Papavasileiou,P., Peters,O. and Rajewsky,N. (2015) Identification and characterization of circular RNAs as a new class of putative biomarkers in human blood. PLoS One, 10, 1–13.

22. Wang,Y., Yu,X., Luo,S. and Han,H. (2015) Comprehensive circular RNA profiling reveals that circular RNA100783 is involved in chronic CD28-associated CD8(+)T cell ageing. Immun. Ageing, 12, 17.

23. Rybak-Wolf,A., Stottmeister,C., Glažar,P., Jens,M., Pino,N., Giusti,S., Hanan,M., Behm,M., Bartok,O., Ashwal-Fluss,R., et al. (2015) Circular RNAs in the Mammalian Brain Are Highly Abundant, Conserved, and Dynamically Expressed. Mol. Cell, 58, 870–885.

24. Corces,M.R., Buenrostro,J.D., Wu,B., Greenside,P.G., Chan,S.M., Koenig,J.L., Snyder,M.P., Pritchard,J.K., Kundaje,A., Greenleaf,W.J., et al. (2016) Lineage-specific and single-cell chromatin accessibility charts human hematopoiesis and leukemia evolution. Nat. Genet., 48, 1193–1203.

25. Kissopoulou,A., Jonasson,J., Lindahl,T.L. and Osman,A. (2013) Next generation sequencing analysis of human platelet polyA+ mRNAs and rRNA-depleted total RNA. PLoS One, 8.

26. Alhasan,A.A., Izuogu,O.G., Al-Balool,H.H., Steyn,J.S., Evans,A., Colzani,M., Ghevaert,C., Mountford,J.C., Marenah,L., Elliott,D.J., et al. (2016) Circular RNA enrichment in platelets is a signature of transcriptome degradation. Blood, 127, e1–e11.

27. Doss,J.F., Corcoran,D.L., Jima,D.D., Telen,M.J., Dave,S.S. and Chi,J.T. (2015) A comprehensive joint analysis of the long and short RNA transcriptomes of human erythrocytes. BMC Genomics, 16, 1–16.

28. Kornienko,A.E., Dotter,C.P., Guenzl,P.M., Gisslinger,H., Gisslinger,B., Cleary,C., Kralovics,R., Pauler,F.M. and Barlow,D.P. (2016) Long non-coding RNAs display higher natural expression variation than protein-coding genes in healthy humans. Genome Biol., 17, 1–23.

29. Cheng,J., Metge,F. and Dieterich,C. (2015) DCC – specific identification and quantification of circular RNAs from sequencing data. Bioinformatics, 32, 1–13.

30. Zhang,X.O., Wang,H. Bin, Zhang,Y., Lu,X., Chen,L.L. and Yang,L. (2014) Complementary sequence-mediated exon circularization. Cell, 159, 134–147.

31. Kolde,R. (2012) Pheatmap: pretty heatmaps.

32. Wickham,H. (2009) ggplot2□: Elegant Graphics for Data Analysis Springer New York, New York, NY.

33. Ovchynnikova,E., Aglialoro,F., Bentlage,A.E.H., Vidarsson,G., Salinas,N.D., Von Lindern,M., Tolia,N.H. and Van Den Akker,E. (2017) DARC extracellular domain remodeling in maturating reticulocytes explains Plasmodium vivax tropism. Blood, 130, 1441–1444.

34. Ye,J., Coulouris,G., Zaretskaya,I., Cutcutache,I., Rozen,S. and Madden,T.L. (2012) Primer-BLAST: a tool to design target-specific primers for polymerase chain reaction. BMC Bioinformatics, 13, 134.

35. Zeng,X., Lin,W., Guo,M. and Zou,Q. (2017) A comprehensive overview and evaluation of circular RNA detection tools. PLoS Comput. Biol., 13, 1–21.

36. Glažar,P., Papavasileiou,P. and Rajewsky,N. (2014) circBase: a database for circular RNAs. RNA, 20, 1666–1670.

37. Liu,Y.C., Li,J.R., Sun,C.H., Andrews,E., Chao,R.F., Lin,F.M., Weng,S.L., Hsu,S. Da, Huang,C.C., Cheng,C., et al. (2016) CircNet: A database of circular RNAs derived from transcriptome sequencing data. Nucleic Acids Res., 44, D209–D215.

38. Matera,A.G. and Wang,Z. (2014) A day in the life of the spliceosome. Nat. Rev. Mol. Cell Biol., 15, 108–121.

39. Pimentel,H., Parra,M., Gee,S., Ghanem,D., An,X., Li,J., Mohandas,N., Pachter,L. and Conboy,J.G. (2014) A dynamic alternative splicing program regulates gene expression during terminal erythropoiesis. Nucleic Acids Res., 42, 4031–4042.

40. Rifkind,J.M. and Nagababu,E. (2013) Hemoglobin Redox Reactions and Red Blood Cell Aging. Antioxid. Redox Signal., 18, 2274–2283.

41. Zheng,Q., Bao,C., Guo,W., Li,S., Chen,J., Chen,B., Luo,Y., Lyu,D., Li,Y., Shi,G., et al. (2016) Circular RNA profiling reveals an abundant circHIPK3 that regulates cell growth by sponging multiple miRNAs. Nat. Commun., 7, 1–13.

42. Kelly,S., Greenman,C., Cook,P.R. and Papantonis,A. (2015) Exon Skipping Is Correlated with Exon Circularization. J. Mol. Biol., 427, 2414–2417.

43. Preußer,C., Hung,L.-H., Schneider,T., Schreiner,S., Hardt,M., Moebus,A., Santoso,S. and Bindereif,A. (2018) Selective release of circRNAs in platelet-derived extracellular vesicles. J. Extracell. Vesicles, 7, 1424473.

44. Greene,J., Baird,A.-M., Brady,L., Lim,M., Gray,S.G., McDermott,R. and Finn,S.P. (2017) Circular RNAs: Biogenesis, Function and Role in Human Diseases. Front. Mol. Biosci., 4, 1–11.

45. Hansen,T.B., Jensen,T.I., Clausen,B.H., Bramsen,J.B., Finsen,B., Damgaard,C.K. and Kjems,J. (2013) Natural RNA circles function as efficient microRNA sponges. Nature, 495, 384–388.

46. Jeck,W.R. and Sharpless,N.E. (2014) Detecting and characterizing circular RNAs. Nat. Biotechnol., 32, 453–461.

47. Yang,Y., Fan,X., Mao,M., Song,X., Wu,P., Zhang,Y., Jin,Y., Yang,Y., Chen,L.-L., Wang,Y., et al. (2017) Extensive translation of circular RNAs driven by N6-methyladenosine. Cell Res., 27, 626–641.

48. Kristensen,L.S., Hansen,T.B., Venø,M.T. and Kjems,J. (2018) Circular RNAs in cancer: Opportunities and challenges in the field. Oncogene, 37, 555–565.

